# Nanoscale dynamics of cholesterol in the cell membrane

**DOI:** 10.1101/644005

**Authors:** Kerstin Pinkwart, Falk Schneider, Martyna Lukoseviciute, Tatjana Sauka-Spengler, Edward Lyman, Christian Eggeling, Erdinc Sezgin

## Abstract

Cholesterol constitutes approximately 30-40% of the mammalian plasma membrane — a larger fraction than any other single component. It is a major player in numerous signalling processes as well as molecular membrane architecture. However, our knowledge on dynamics of cholesterol in the plasma membrane is limited which restricts our understanding of the mechanisms regulating its involvement in cell signalling. Here, advanced fluorescence imaging and spectroscopy approaches were applied on *in vitro* (model membranes) and *in vivo* (live cells and embryos) membranes to systematically study the nanoscale dynamics of cholesterol in biological membranes. The results show that cholesterol diffuses faster than phospholipids in live membranes, but not in model membranes. The data indicate that diffusion of cholesterol and phospholipids is not correlated with membrane domain partitioning. Instead, our data show that the fast diffusion of cholesterol is due to its nanoscale interactions and localization in the membrane.

## Introduction

Cholesterol plays a pivotal role in eukaryotic cellular membranes both structurally and functionally^1-3^. Besides its involvement in signaling ^4-8^, it considerably shapes the molecular architecture of the plasma membrane^9, 10^. However, due to the challenge of performing high resolution measurements on small and fast molecules in cells, molecular-level knowledge is limited regarding specific mechanisms involving cholesterol. To better understand how cholesterol influences cellular signaling, it is essential to directly observe the dynamics of cholesterol in the plasma membrane.

Fluorescence microscopy and spectroscopy are powerful tools for studying molecular dynamics in living cells^11^. Visualization of cellular cholesterol with these approaches is usually achieved using fluorescently-labeled cholesterol analogs^11^. These analogues, however, may not reflect the native behavior of cholesterol because the cholesterol and the fluorescent tag are of similar size — a design consideration for all fluorescent lipid analogues. One important feature of robust lipid analogs is that they partition into membrane regions like their unlabeled counterparts. Ordered membranes (or nanodomains) are enriched with saturated lipids, some of which (such as sphingomyelin) have preferential cholesterol binding^12^. Therefore, the cholesterol analogs that are enriched in the ordered plasma membrane domains are considered to be reliable mimics of native cholesterol^2, 9^. Another important criterion for the reliability of fluorescent lipid analogues is proper cellular trafficking into subcellular structures. Bodipy-labelled cholesterol, for instance, has shown a great potential to mimic cholesterol in terms of partitioning, dynamics and cellular localization^13-17^. Despite its ordered domain partitioning, interestingly, recent reports suggested an order of magnitude faster diffusion for cholesterol compared to phospholipids^18, 19^. This finding brings a conundrum; how can cholesterol move faster in the cell membrane while being enriched in more ordered (packed) membrane environments? To answer this, studies aiming at revealing the nanoscale cholesterol diffusion in the cell membranes and its determinants are needed.

Nanoscale behavior of molecules is not readily accessible with conventional methodologies. The investigation of nanoscale dynamics of specific molecules in the membrane has been hampered by the complex, heterogeneous, and hierarchical structure of the cell membrane. Moreover, nanoscale interactions occur at very fast temporal and small spatial scales, which presents additional experimental challenges. Therefore, a complete understanding of nanoscale molecular behavior requires both advanced imaging techniques and well-defined model membrane systems. Here, we applied advanced imaging and spectroscopy approaches on *in vitro* (model membranes) and *in vivo* (live cells and embryos) membranes to elucidate the nanoscale diffusion dynamics of cholesterol in biological membranes. We compared the diffusion of cholesterol with that of phospholipids as well as of cholesterol analogues carrying acyl chains. The experimental measurements were complemented by *in silico* Monte Carlo and Molecular Dynamics simulations. The data show that cholesterol diffuses faster than phospholipids in cellular membranes, particularly in live membranes. Furthermore, membrane domain partitioning does not influence the nanoscale diffusion, whereas nanoscale interactions and localization in the plasma membrane significantly influence the diffusional dynamics of cholesterol.

## Results and Discussion

To understand cholesterol dynamics at the nanoscale in biomembranes, we first compared the diffusion dynamics between various cholesterol analogues and phospholipid analogues (Figure 1). Fluorescent lipid labelling is generally a challenging issue due to the comparable sizes of the lipids and the fluorescent tags. It is particularly a major problem for cholesterol labelling due to its comparably smaller size^20^. A few intrinsically fluorescent analogues ^21-23^ and Bodipy-labelled (trademarked as Topfluor (TF)) cholesterol were shown to mimic cholesterol behaviour in model and cell membranes^13, 14, 16, 24, 25^. Similarly a few NBD-labelled cholesterol were reported to mimic certain properties of cholesterol^26, 27^. Recently, cholesterol was also tagged with different fluorophores (e.g. Abberior Star Red (AbStarRed)) via a PEG linker in between the fluorophore and sterol moiety^28^ to avoid the fluorophore interaction with the membrane^29^. Here, we employed TF-Chol and AbStarRed-PEG-Chol as cholesterol analogues. We also used TF-Chol attached to a saturated (16:0) and an unsaturated (18:2) acyl chain to study whether the acyl chains make cholesterol behave similar to phospholipids in the membrane. In addition, we used phosphatidylethanolamine (PE) and sphingomyelin (SM) tagged with Topfluor (Figure 1) to directly compare the cholesterol diffusion with other lipid species.

**Figure 1.**
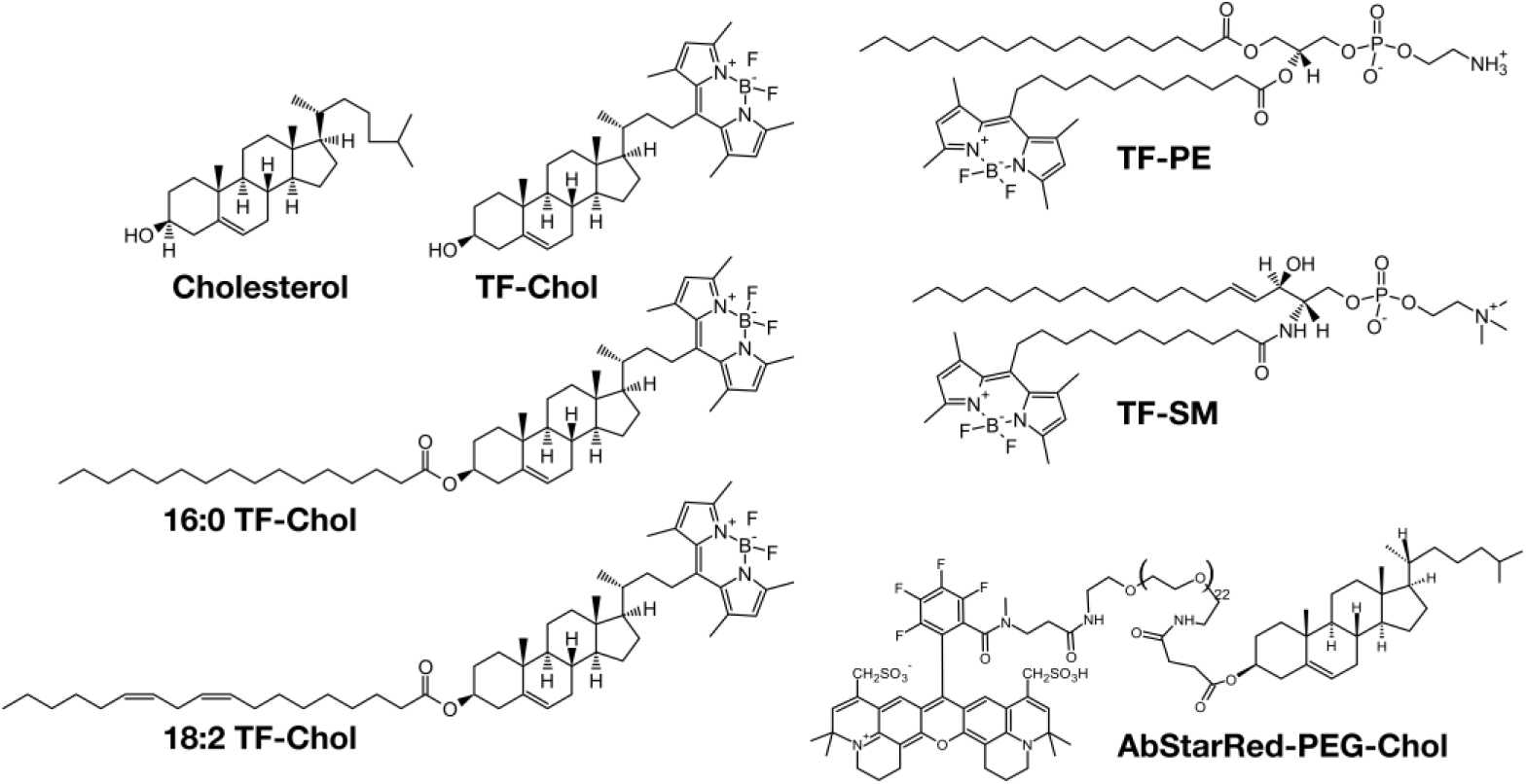
Structures of fluorescent lipid analogues used in this study.

### Diffusion of cholesterol compared to phospholipids in model membranes

Previous reports suggested fast diffusion of TF-Chol (≈ 3 µm^2^/s) in the plasma membrane of live cells^18, 19^. To tackle how cholesterol moves compared to phospho- and sphingolipids in the cell membrane systematically, we used simple artificial model membranes (giant unilamellar vesicles; GUVs), cell-derived vesicles (giant plasma membrane vesicles; GPMVs), live mammalian cells and zebrafish embryos. First, we compared whether an acyl chain attachment changes the cholesterol mobility. To this end, we compared the diffusion of TF-Chol with TF-Chol bearing either a saturated or an unsaturated acyl chains (16:0 TF-Chol and 18:2 TF-Chol, respectively) (Figure 1). We stained the synthetic GUVs and cell-derived GPMVs with low amount of the fluorescent analogues and Cell Mask deep red for membrane visualization (Figure 2A, B). Then, we performed fluorescence correlation spectroscopy (FCS) to obtain the diffusion coefficients for each fluorescent analogue. In GUVs, acyl-chain carrying cholesterol analogues did not show notably different diffusion (D_16:0_ = 10.0±0.9 µm^2^/s; D_18:2_ = 10.3±0.8 µm^2^/s) compared to TF-Chol (D_TF-Chol_ = 10.5±1.2 µm^2^/s) (Figure 2C). TF-PE and TF-SM showed slightly slower (≈1.2 times) diffusion with diffusion coefficients of 8.9±1.4 µm^2^/s and 8.8±1.1 µm^2^/s, respectively (Figure 2C) compared to TF-Chol. In GPMVs isolated from Chinese Hamster Ovary (CHO) cells, addition of an acyl chain only slightly changed the diffusion coefficient of the TF-Chol analogues (Figure 2D). There was, however, a clear difference between TF-Chol and TF-PE or TF-SM (Figure 2D; D_TF-Chol_ = 4.9±0.7 µm^2^/s compared to D_TF-PE_ = 4.0±0.8 µm^2^/s, D_TF-SM_ = 3.9±0.8 µm^2^/s). The lipid analogues were slower in GPMVs compared to GUVs and in both model systems TF-Chol diffusion was ≈1.2 times faster than TF-PE or TF-SM. This difference was small but reproducible (Supplementary Figure S1).

**Figure 2.**
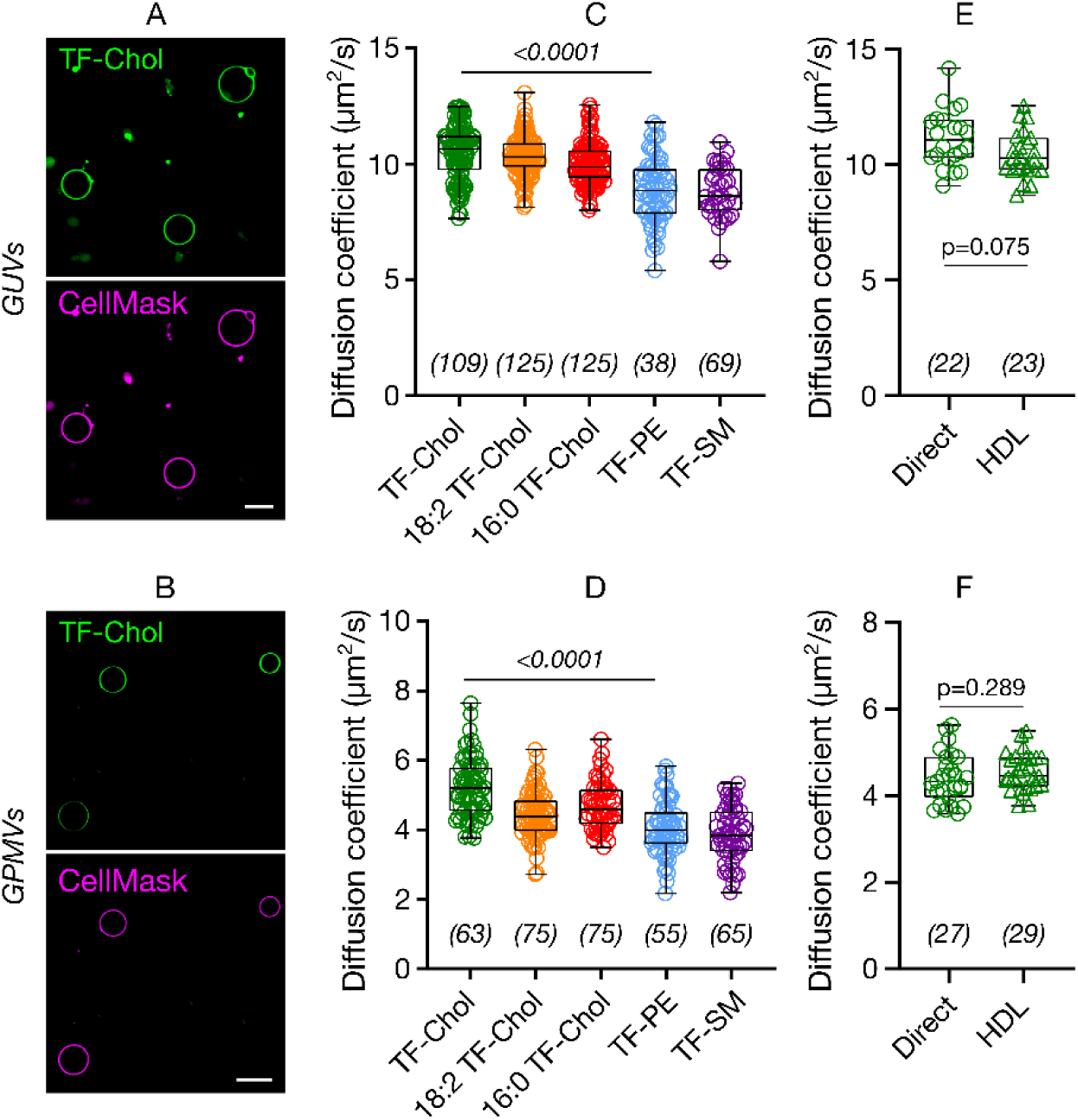
Diffusion of cholesterol compared to phospholipids in model membranes. Confocal images of A) GUVs and B) GPMVs labelled with TF-Chol (green) and CellMask (magenta) which labels the membrane. Scale bars are 10 µm. Diffusion of cholesterol and phospholipid analogues in C) GUVs and D) GPMVs. TF-Chol diffusion in E) GUVs, F) GPMVs with direct labelling or via HDL particles. Data are shown as box-and-whisker plot showing median, first and third quartiles, and all the data points. Number of data points are indicated on the graphs.

Cell membranes can be labelled with cholesterol analogues via different approaches such as direct labelling through incubation of the sample with pure TF-Chol or with high and low density lipoproteins (HDL, LDL) particles carrying fluorescent cholesterol^16, 30, 31^. The latter is of great efficiency since these particles primarily manage cholesterol transfer in the body. It should be taken into account that the delivery methods may change how the cholesterol is incorporated into the cell membranes that might affect its subsequent diffusion. To this end, we also measured the diffusion of TF-Chol in GUVs and GPMVs where vesicles are labelled either directly by adding the TF-Chol to the vesicle suspension or via HDL particles carrying TF-Chol. Both in GUVs and in GPMVs, cholesterol diffusion via direct labelling and via HDL particles yielded the same diffusion coefficients (Figure 2E, F).

### Diffusion of cholesterol compared to phospholipids in live membranes

Next, we investigated diffusion of cholesterol in live membranes. We tested the diffusion of cholesterol, phospholipid and sphingomyelin analogues in CHO cells as an *in cellulo* model for mammalian cells (Figure 3A) and in zebrafish embryos as an *in vivo* model (Figure 3B, see Supplementary Figure S2 for details of FCS measurements). In both of these systems we observed markedly faster diffusion of TF-Chol compared to TF-PE or TF-SM (Figure 3C). In CHO cells, the diffusion coefficient was 2.4 µm^2^/s for TF-Chol, 1.4 µm^2^/s for TF-PE and 1.1 µm^2^/s for TF-SM. In zebrafish embryos, diffusion coefficient was 2.8 µm^2^/s for TF-Chol, 1.1 µm^2^/s for TF-PE and 1.5 µm^2^/s for TF-SM. Overall, we observed ≈2-fold faster diffusion of TF-Chol compared to TF-labelled phospholipid and sphingolipid analogues *in vivo*. In model membranes this difference was ≈1.2 times which suggests active processes in living membranes lead to slower diffusion of phospholipids and sphingolipids compared to cholesterol.

**Figure 3.**
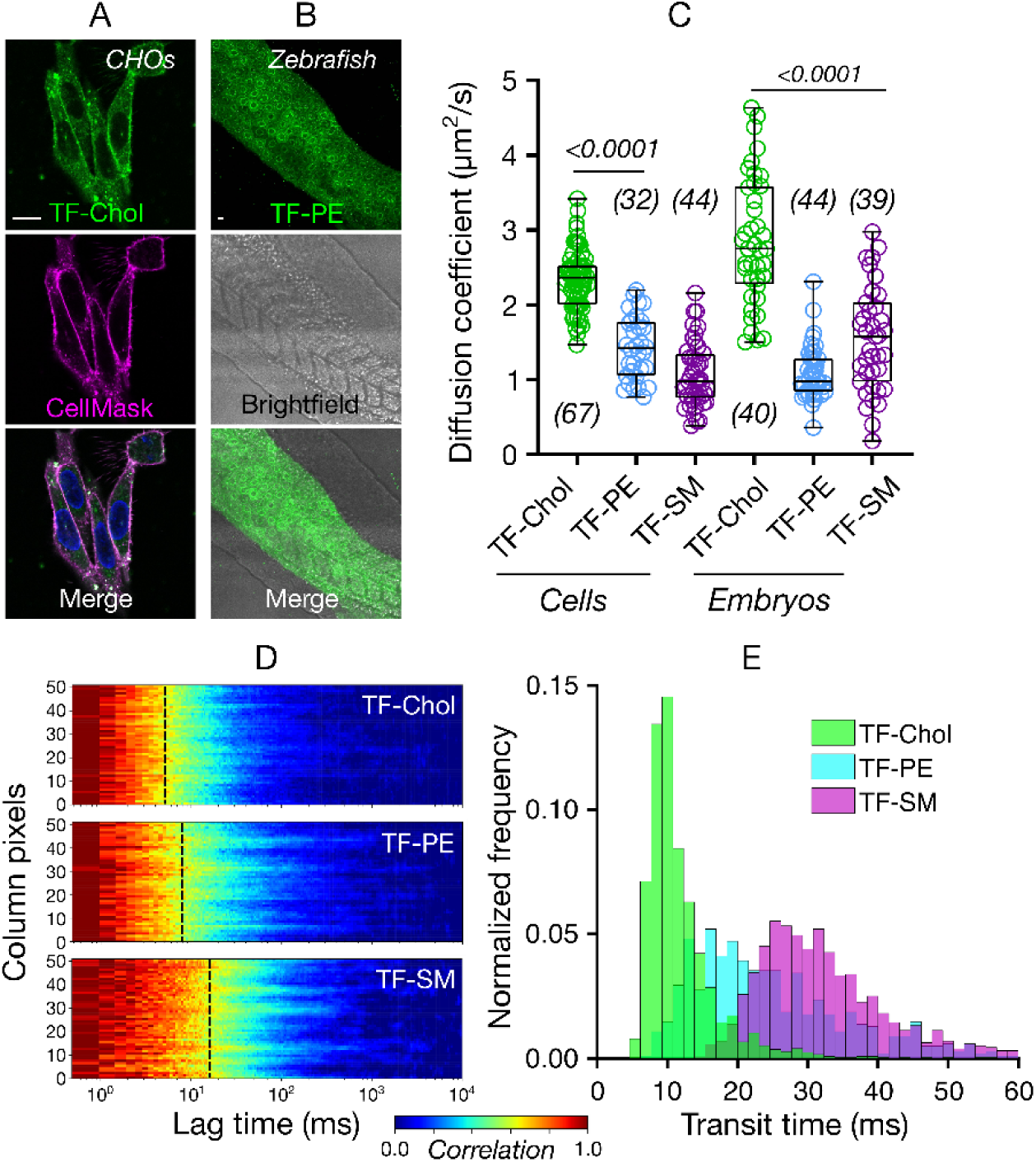
Diffusion of cholesterol compared to phospholipids *in cellulo* and *in vivo*. Confocal images of A) CHO cells B) zebrafish embryos (24 hours post fertilization) labelled with fluorescent lipid analogues (green) and Cell Mask Deep Red (magenta). Scale bars are 10 µm. C) Diffusion coefficient of the fluorescent lipid analogues in CHO cells and zebrafish embryos. D) Representative sFCS carpets for fluorescent lipid analogues in CHO cells. E) Histogram of transit diffusion times of the fluorescent lipid analogues obtained from the sFCS data. Data are shown as box-and-whisker plot showing median, first and third quartiles, and maximum and minimum values. Number of data points are indicated on the graphs.

While the diffusion data obtained with point FCS is extremely informative, it is limited to a diffraction-limited spot and may overlook spatial heterogeneity. The remedy for this is scanning FCS (sFCS) which reports on molecular diffusion along a line (≈5 µm long, ≈50 pixels), thus captures the spatial diffusional heterogeneity in the plasma membrane and better probes the nanoscale dynamics^32^. Moreover, sFCS yields multiple curves (as many as the number of pixels) per measurement which increases the statistical accuracy of the measurements^28^. We, thus, performed sFCS to study the diffusion of fluorescent lipid analogues in the cell membrane with more accurate spatial sampling. sFCS carpets (which are color-coded FCS curves, red showing the maximum correlation, blue showing the minimum correlation) on live CHO cells show clear fast diffusion of TF-Chol compared to TF-PE and TF-SM (yellow parts corresponds to the transit diffusion time, Figure 3D). Histogram of the transit times obtained from the sFCS data confirms the fast diffusion of TF-Chol, slower diffusion of TF-PE and slowest diffusion of TF-SM (Figure 3E).

### Diffusion of fluorescent lipid analogues vs. their membrane domain partitioning

We next set out to determine which aspects of living membranes cause faster diffusion of cholesterol in cells or, more precisely, slower diffusion of phospholipid and sphingolipid analogues. One explanation is differential partitioning into membrane environments of different fluidity and packing. Molecules diffuse more slowly in ordered plasma membrane environments enriched with saturated lipids compared to disordered membranes enriched with unsaturated lipids^33^. Coexistence of ordered and disordered domains is observed in model membranes depending on composition and temperature. The most physiological systems where such macroscopic phase separation is observed are giant plasma membrane vesicles (GPMVs) derived from live cells^34, 35^. Therefore, to test whether the discrepancy between the diffusion of the analogues we observe can be attributed to heterogeneous partitioning in the plasma membrane, we correlated partitioning in phase-separated GPMVs with diffusion in cell membranes. TF-Chol and TF-SM partitioned into the ordered domains (≈60%) while others predominantly partitioned into the disordered domain (Figure 4A, B). When we compared the ordered domain partitioning with the diffusion properties, we observed no correlation between the diffusion and the partitioning (Figure 4C). Therefore, we conclude that the slow diffusion of TF-PE and TF-SM is not related to their partitioning into membrane domains which suggests that the domain incorporation may not have a drastic effect on probe diffusion.

**Figure 4.**
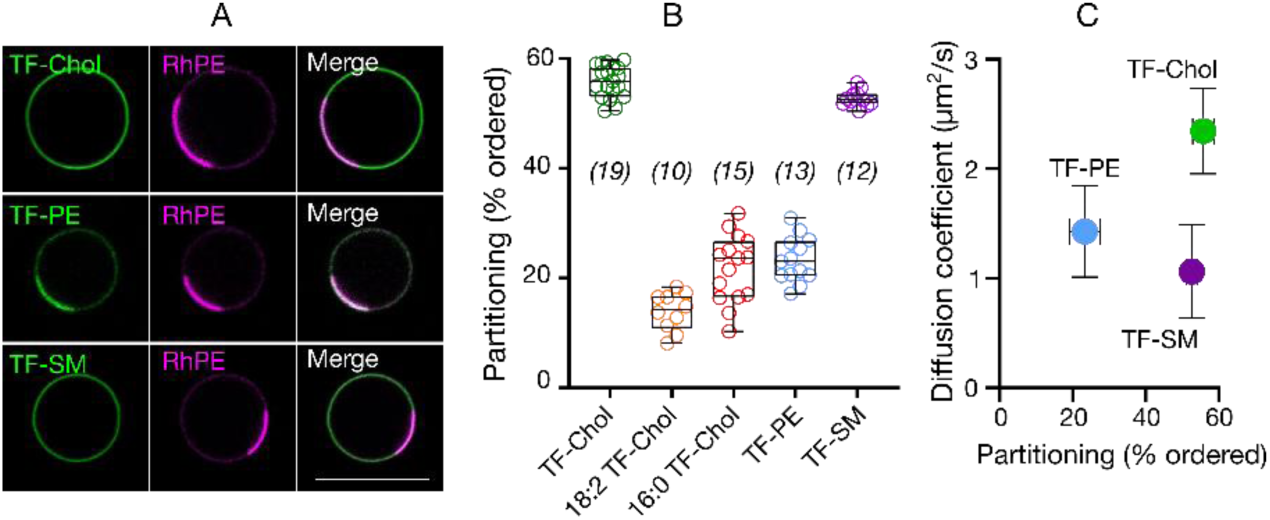
Relationship between diffusion in cells and analogue partitioning in phase separated GPMVs. A) Confocal images of TF-Chol, TF-PE and TF-SM partitioning in phase separated GPMVs with Rhodamine-PE (RhPE) as disordered phase marker. Scale bar is 10 µm. B) Quantification of ordered domain partitioning of fluorescent lipid analogues from the confocal images. C) Correlation between ordered domain partitioning in GPMVs and diffusion of fluorescent lipid analogues in cells. Data are shown as box-and-whisker plot showing median, first and third quartiles, and all the data values. Number of data points are indicated on the graphs.

### Diffusion of fluorescent lipid analogues vs. their nanoscale interactions

Another explanation for the slow diffusion of TF-PE and TF-SM relative to cholesterol in live cell membranes are nanoscale hindrances to diffusion that affect phospholipids, but not cholesterol. A “hindered diffusion mode” is often reported for various membrane components, but has not been systematically studied for cholesterol. The key to such measurements is to determine how the apparent diffusion coefficient of molecules changes with the size of the observation spot^36^. For a molecule undergoing Brownian (or free) diffusion, the apparent diffusion coefficient is independent of the size of the observation spot, while for hindered diffusion, the diffusion coefficient varies depending on spot size. The qualitative dependence of the effective diffusion constant on observation volume depends on the underlying diffusion law. It decreases with decreasing observation spot size when molecules are transiently immobilized or trapped or incorporated into slow-moving molecular complexes with sizes below the diffraction limit ^37, 38^. Conversely, it will increase with decreasing spot size when molecules undergo hop or compartmentalized diffusion in a meshwork-like pattern^39-41^.

To discern the role of hindrances in the plasma membrane on the slower diffusion of PE and SM analogues (and faster diffusion of cholesterol), we first simulated the diffusion of cholesterol and phospholipids *in silico* by molecular dynamics simulations in a ternary mixture of DPPC/DOPC/Chol, with the composition chosen to ensure a liquid-disordered membrane. Cholesterol diffusion was slightly faster (≈1.15 times) than phospholipids (Figure 5A), which is in accordance with the experimental model membrane data (≈1.2 times, Figure 2A-D). (Note that the absolute diffusion coefficients vary between *in silico* vs the model membrane experiments due to a finite-size effect^42^.

**Figure 5.**
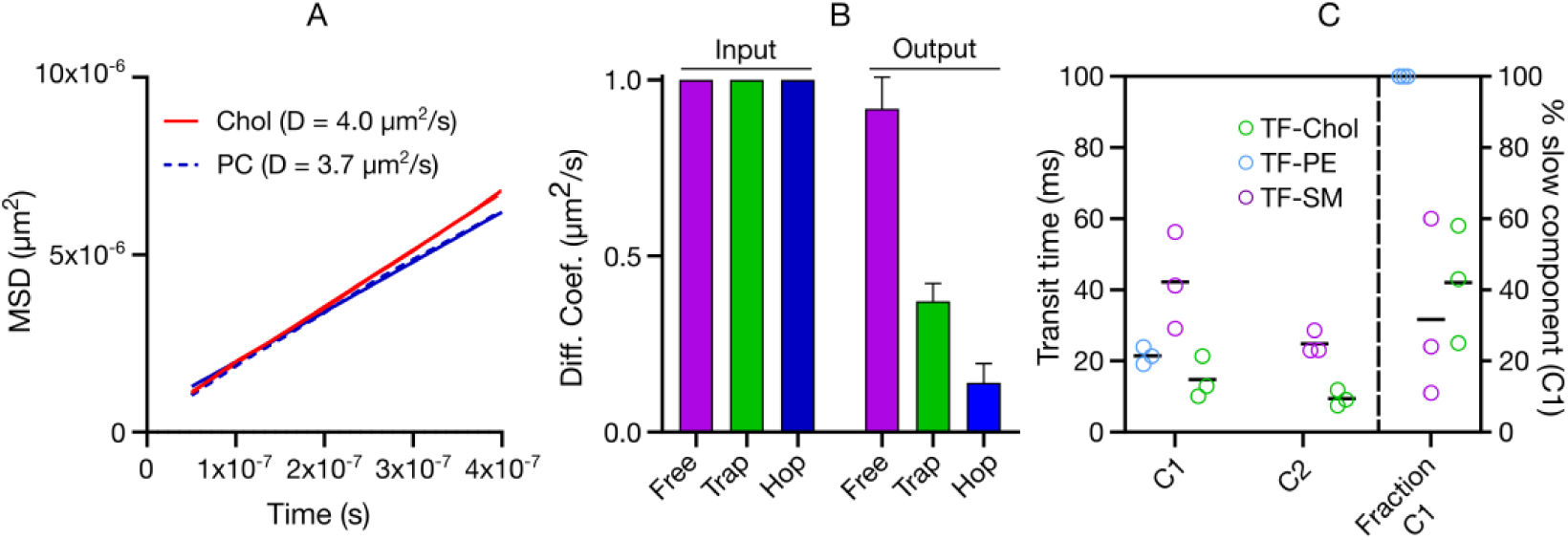
Diffusion mode of fluorescent lipid analogues. A) All-atom molecular dynamics simulation of cholesterol in liquid-disordered DPPC/DOPC/Chol membrane comparing cholesterol with phospholipid diffusion. B) Monte Carlo simulations of free, hop and trapped diffusion probed by FCS showing how the hindrances affect the molecular diffusion. “Input” are the starting diffusion coefficients and “Output” are the diffusion coefficients when free diffusion or hindered diffusion (trapping or hopping) is employed in the simulations. C) Statistical analysis of sFCS data showing that TF-PE can be fit with one component while TF-SM and TF-Chol have two diffusional components; C1:slow component and C2:fast component. Mean and standard deviation of at least 10 independent simulations per condition is shown in panel B. In panel C, each data point shows a repetition of the sFCS measurement and each point consists of averaged data of >10 cells and >1000 curves. Black lines show the mean of the three repetitions.

We also simulated how the diffusion coefficients change when different hindrances are introduced (Figure 5B) using Monte Carlo as described in Methods section. For these simulations, we initialized the simulations with molecules having a diffusion coefficient of ≈1 µm^2^/s. We simulated molecules undergoing free, hop and trapped diffusion modes. Although the simulations for all molecules were initialised with the same diffusion coefficients (that is, the microscopic diffusion constant in the absence of hindrances was 1 µm^2^/s), the diffusion coefficients of the molecules undergoing the hop and trapped diffusion yielded lower values showing that these hindrances actively slows down the diffusion of the molecules.

Cholesterol has previously been investigated with STED-FCS^15^; the data were consistent with free diffusion^19^. However, as discussed above, this measurement is not sensitive to spatial heterogeneity. We therefore performed scanning FCS statistical analysis to investigate the diffusion modes of TF-labelled analogues with high spatiotemporal and statistical accuracy. In this analysis, we generate histograms from all the FCS transit times and fit these with a log-normal distribution which yields two parameters (mean µ and standard deviation σ) ^43^. For a freely diffusing molecule, µ is closely related to the median transit time of the population (*e*^*µ*^ = median transit time) and σ accounts for the skew and spread of the distribution caused by the slow sampling in sFCS (Supplementary Figure S3). For molecules undergoing free diffusion, a simple lognormal fit represents the data well, unlike for molecules undergoing hindrance where a lognormal fit does not fit the data. In this case, a two component (double lognormal) fit is required to describe the data, where the second component accounts for a fraction (1-B) of molecules with slower diffusion due to nanoscale interactions. We use the Bayesian Information Criterion for unbiased model selection^43-45^ to decide between single and double log-normal distribution. Over the length scales accessible with this methodology, the data showed that TF-PE exhibits free diffusion with a single component with transit diffusion time (*e*^*µ*^) of 21 ± 2 ms (Figure 5C). TF-SM showed two different diffusional components; slow component C1 with a diffusion time scale of *e*^*µ*^ ≈ 42 ± 13 ms and a fast component C2 with *e*^*µ*^ ≈ 25±3 ms) (Figure 5C, Supplementary Figure S3) which is a usual manifestation of trapped diffusion (as observed before with statistical analysis for Atto647N-SM^43^). The fast component (C2) of TF-SM is only slightly slower than TF-PE which is in line with the model membrane data (see Figure 2). The slow component (C1) was ≈2 times slower than TF-PE. Surprisingly, TF-Chol also showed two diffusional components (Figure 5C). One component (*e*^*µ*^ ≈ 15 ± 6 ms) was slightly faster than TF-PE (≈1.4 times), a difference similar to what we observed in model membranes (see Figure 2). The fast component (*e*^*µ*^ ≈ 9 ± 2 ms) was significantly faster (≈2.3 times) than TF-PE.

### Diffusion of fluorescent lipid analogues vs. their localization

Two-component diffusion of TF-SM can be explained by its trapping behaviour which was observed previously with various methodologies and different SM analogues^38, 46-48^. However, the two-component diffusion of cholesterol cannot be explained by simple trapping since one component is faster but not slower than the membrane diffusion of TF-PE. Moreover, previously TF-Chol has been reported to exhibit free diffusion^18, 19^. It is postulated that the location of cholesterol (e.g. in different leaflet or its location relative to the membrane interface) in the membrane can drastically influence its diffusion^19^. Leaflets in the plasma membrane are asymmetric; glycosphingolipids, sphingomyelin and phosphatidylcholine are predominantly in the outer leaflet while phosphatidylserine and phosphatidylethanolamine are mostly in the inner leaflet^49^. This asymmetric composition creates a difference in the membrane organisation thus different microenvironment in the inner vs outer leaflets. Therefore, the fast diffusion of cholesterol compared to SM and PE in live cells may be due to their different leaflet preferences. There are contradictory reports on the leaflet preference of cholesterol; whereas some reports suggest outer leaflet enrichment of cholesterol^50^, some recent reports contradict this hypothesis^51, 52^ while others suggest more dynamic partitioning of cholesterol between leaflets^53^. As inner and outer leaflets have different ordering (presumably with the inner leaflet being more fluid^54^), fast diffusion of cholesterol may be caused by differential partitioning of cholesterol. It was also previously speculated that alignment of cholesterol relative to the membrane plane alters its diffusion^19^. Interestingly, recent studies suggested cholesterol localization in the mid-plane of the bilayer (between the leaflets)^55^ which would heavily influence the diffusion of cholesterol. Therefore, we set out to address whether cholesterol localization in the membrane has an influence on its diffusion. First, we tested whether TF-Chol has different geometry inside the bilayer compared to TF-PE and TF-SM which would be a strong indication of different localization. To this end, we used single molecule fluctuation analysis of sFCS data. Fluorophore alignment with respect to the membrane plane varies its extinction coefficient. The fluorophores are more efficiently excited (i.e., higher brightness) when the dipole moment of the fluorophore is parallel to the polarization of the laser. When we compared the brightness (photons per single particle) of the TF-PE, TF-SM and TF-Chol in live cell membranes, we found out that TF-Chol brightness was significantly higher than TF-PE and TF-SM which suggest that TF-Chol geometry, and presumably localization in the membrane is different (Figure 6A). To further test whether membrane localization has impact on diffusion, we used a cholesterol analogue, Abberior Star Red-PEG-Chol (AbStR-Chol) that cannot flip/flop in the membrane due to the PEG linker between the cholesterol and the dye and is located exclusively in the outer leaflet^29, 56^. We compared its diffusion with Abberior Star Red-PEG-PE and Abberior Star Red-SM which are also located in the outer leaflet. They stained the plasma membrane with no notable internal signal which suggests that they are not flipped to the inner part of the cell (Figure 6B). The diffusion of AbStR-Chol, AbStR-PE and AbStR-SM was approximately the same (≈1.0 µm^2^/s) and significantly slower than TF-Chol (≈2.4 µm^2^/s) indicating that cholesterol diffusion is as slow as phospholipid diffusion when located in the outer leaflet. Overall, this data suggests that cholesterol localization in the membrane is more complex than phospholipids and this localization influences its diffusion.

**Figure 6.**
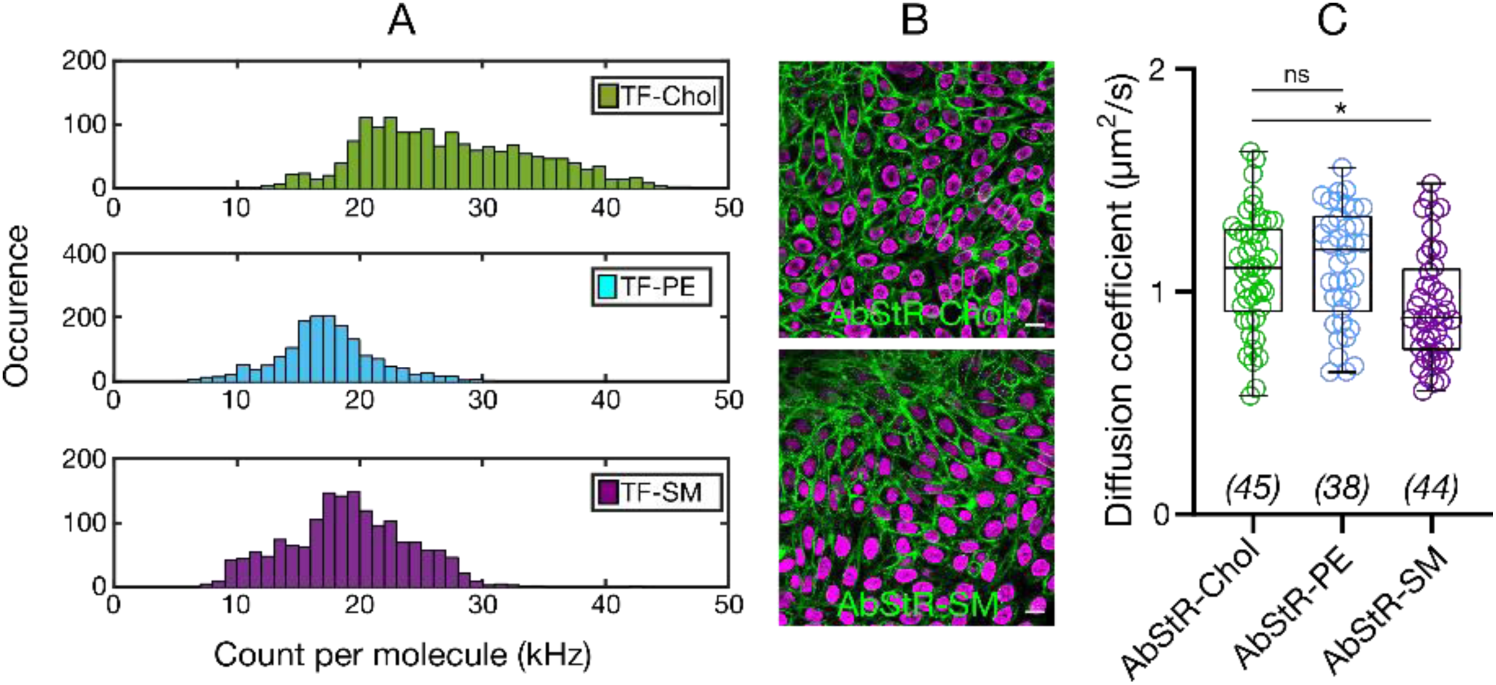
A) Count per molecule (cpm) histograms determined from sFCS data for TF-Chol, TF-PE and TF-SM in CHO cells showing higher cpm values for TF-Chol (>1000 curves). B) Confocal images of CHO cells labelled with non-flipping Abberior Star Red-PEG-Chol and Abberior Star Red-SM. Scale bars are 10 µm. C) Diffusion coefficient of Abberior Star Red-PEG-Chol, Abberior Star Red-PEG-PE and Abberior Star Red-SM in CHO cells. Data are shown as box-and-whisker plot showing median, first and third quartiles, and all the data values. Number of data points are indicated on the graphs.

## Conclusion

Cholesterol is one of the most important membrane components, involved in cellular membrane structure and signaling. Building more quantitative models for the role of cholesterol in signalling demands a better understanding of its lateral diffusion. Here, we investigated the diffusion of fluorescent cholesterol analogues in both model and cellular membranes using advanced imaging and spectroscopy tools as well as molecular simulations. We found that cholesterol moves only slightly faster (≈1.2 times) than phospholipids and sphingolipids in model membranes that are thermodynamically in equilibrium. However, in live cells and embryos, it diffused significantly faster (≈2 times) than phospholipids and sphingolipids. We found out that that the slower diffusion of phospho- and sphingolipids (and thus faster diffusion of cholesterol) is not due to membrane domain partitioning. The fluorescent cholesterol analogue TF-Chol showed two diffusional components in live cells; one being similar to fluorescent PE analogue TF-PE and the other one being faster. On the other hand, TF-SM also showed two diffusional components; one being similar to TF-PE while the other one being slower. This slow diffusional component of TF-SM is indication of trapping^38^. Brightness analysis revealed that the fast diffusional component of TF-Chol in contrast was due to heterogeneous localization/positioning of cholesterol within the membrane, e.g. asymmetric partitioning between the inner and outer leaflets of the plasma membrane and localization into the mid-plane of the bilayer proposed^55^. This is supported by the similar diffusion of cholesterol and phospholipid analogues when they are localised to the outer leaflet of the cell membrane.

Our results suggest an asymmetric localisation of cholesterol in the membrane. These findings may be used to shed light on membrane asymmetry and heterogeneity. Moreover, our findings are crucial for membrane biology and naturally create new questions; how can cholesterol contribute to high order membrane domain formation with sphingomyelin if their diffusion dynamics are so different? Are the fast and slow components of SM and Chol different and related to recently reported different forms of SM^57^? Do interactions occur between the fast SM and slow cholesterol which have similar diffusion dynamics? What is the role of fast-moving cholesterol? And finally how can cholesterol contribute to cellular signalling with such fast diffusion dynamics? Many proteins have been proposed to have cholesterol binding motif^3, 6, 8, 58, 59^. It will also be essential to address how consensus cholesterol-binding motifs have been found in different membrane leaflets and whether it is related to cholesterol leaflet preference^60^.

It is important to note that, here we use TF-Chol, a fluorescent cholesterol analogue that has been proven reliable for many aspects. However, there is still risk of artefacts induced by the fluorescent probe, thus further studies with label-free technologies will be useful to address these vital question in cell biology. Further to that, cholesterol localization and flipping may be critical for domain formation as suggested recently^61, 62^. Thus, answers to these questions will also contribute to the efforts to understand the functional membrane heterogeneity. Our present measurement and experimental approaches give further guides to solving such long-standing questions.

## Experimental

### Lipids and fluorescent lipid analogs

We purchased 23-(dipyrrometheneboron difluoride)-24-norcholesterol (Topfluor Cholesterol; TF-Chol), 23-(dipyrrometheneboron difluoride)-24-norcholesteryl palmitate (16:0 TF-Chol), 23-(dipyrrometheneboron difluoride)-24-norcholesteryl linoleate (18:2 TF-Chol) 1,2-Dioleoyl-sn-glycero-3-phosphocholine (DOPC), Topfluor-sphingomyelin (TF-SM), Rhodamine PE and Topfluor-phosphoethanolamine (TF-PE) from Avanti Polar Lipids. Abberior Star Red-PEG-Cholesterol was obtained from Abberior. Cell Mask and NucBlue were obtained from Thermofisher.

### Preparation of GUVs

GUVs were prepared with electroformation method as previously described^63^. Briefly, a lipid film was formed on a platinum wire from 1 mg/ml lipid mix (DOPC). Then, GUVs were formed in 300 mM sucrose solution at room temperature. 10 Hz, 2 V alternative electric current was used for electroformation.

### Cell culture and zebrafish embryos

CHO cells were maintained in DMEM-F12 medium supplemented with 10% FBS medium and 1% L-glutamine.

For the zebrafish embryos, both females and males of wild-type zebrafish strain were used. Breeding animals were between 3 months old and 2 years old. Zebrafish embryos that were used for the experiments were 24 hours post fertilisation. Animals were handled in accordance to procedures authorized by the UK Home Office in accordance with UK law (Animals [Scientific Procedures] Act 1986) and the recommendations in the Guide for the Care and Use of Laboratory Animals. All vertebrate animal work was performed at the facilities of Oxford University Biomedical Services. Adult fish were maintained as described previously^64^. In brief, adult fish were exposed to 12 hour light – 12 hour dark cycle (8am to 10pm light; 10pm to 8am dark), kept in a closed recirculating system water at 27-28.5°C, fed 3-4 times a day, kept at 5 fish per 1L density. Embryos were staged as described previously^65^.

### Giant Plasma Membrane Vesicles (GPMVs)

GPMVs were prepared as previously described^34^. Briefly, cells seeded out on a 60 mm petri dish (≈70 % confluent) were washed with GPMV buffer (150 mM NaCl, 10 mM Hepes, 2 mM CaCl2, pH 7.4) twice. 2 ml of GPMV buffer was added to the cells. 25 mM PFA and 2 mM DTT (final concentrations) were added in the GPMV buffer. The cells were incubated for 2 h at 37 °C. Then, GPMVs were collected by pipetting out the supernatant. For phase-separated GPMVs, 20 mM DTT was used instead of 2 mM. To observe phase separation, cooling GPMVs to 10 °C may be necessary depending on the cell types.

### Fluorescent Labelling of GUVs, GPMVs, cells and zebrafish embryos

Tips of the pipette tips were cut before handling the GUVs to avoid GUV rupturing due to the shear stress. GUVs and GPMVs were labelled by adding the lipid analogues as well as Cell Mask and rhodamine PE to a final concentration of 10-50 ng/mL. For HDL labelling, 100 µl of vesicle suspension were incubated in 100 ng/ml HDL (gift from Prof. Herbert Stangl) for 30 minutes. Labelled GUVs were placed in the wells of BSA-coated 8-well glass bottom Ibidi chambers which were filled with 250 µL of PBS.

For the cell labeling, the cells were seeded on 25 mm diameter round coverslips (#1.5) in a 6 well plate 2-3 days before the measurements. The fluorescent lipid analogs were first dissolved in DMSO or ethanol with a final concentration of 1 mg/ml. Before the labelling, the cells seeded on glass slides were washed twice with L15 medium to remove the full media. Please note that serum in the media decreases the labelling efficiency, thus it is crucial to wash out all the media from the cells. Later the fluorescent analogs were mixed with L15 medium with 1:1000 ratio (final concentration of 1 µg/ml). The cells were incubated with this suspension for 5-10 minutes at room temperature. After that, the cells were washed with L15 twice followed by imaging in the same medium. Then, confocal microscopy was performed as described below. Labelling cells with fluorescent analogs should be optimized for every cell line by changing the concentration, labelling time and labelling temperature. For nucleus labelling, NucBlue live nucleus staining was done. One drop of NucBlue was added in 1 ml of medium and the cells were incubated in this solution for 15 minutes.

Zebrafish embryos (24 hpf) were incubated in E3 buffer (4.5mM NaCl, 0.18mM KCl, 0.33mM CaCl2•2H20, 0.4mM MgCl2•6H20 in water) for 24 hours and dechorionated manually using forceps. Later, they were mixed with lipid analogs (final concentration of 0.5 µg/ml) for 1 hour at 28 °C followed by another 1 hour of a gentle nutation at room temperature. The embryos were transferred to a 10 cm petri dish filled with fresh E3 solution for washing. Later, the embryos were washed twice with E3 medium and transferred to the Ibidi chambers filled with 250 µL E3 buffer for imaging.

### Confocal microscopy

GUVs were imaged in PBS, GPMVs were imaged in GPMV buffer, cells were imaged in L15 medium and embryos were imaged in E3 buffer. All imaging was done at room temperature (21-23 °C). All imaging was done on glass slides with thickness of 0.17 mm. Samples were imaged with a Zeiss LSM 780 (or 880) confocal microscope in BSA-coated (1 mg/ml for 1 h) 8-well Ibidi glass chambers (#1.5). TF-labeled analogs were excited with 488 nm and emission collected between 505-550 nm. Abberior Star Red-labeled analogs as well as Cell Mask Deep Red were excited with 633 nm and emission collected with 650-700 nm. NucBlue was excited with 405 nm and emission was collected at 420-470 nm. Multi-track mode was used to avoid cross-talk.

### Fluorescence Correlation Spectroscopy (FCS)

FCS on the GUVs and GPMVs were carried out using Zeiss LSM 780 (or 880) microscope, 40X water immersion objective (numerical aperture 1.2) as described before^48^. Briefly, before the measurement, the shape and the size of the focal spot was calibrated using Alexa 488 and Alexa 647 dyes in water in an 8-well glass bottom (#1.5) chamber. To measure the diffusion on the membrane, GUVs and GPMVs were placed into an 8-well glass bottom (#1.5) chamber coated with BSA. The laser spot was focused on the top membrane by maximising the fluorescence intensity. Then, 3-5 curves were obtained for each spot (five seconds each). Cell measurements were performed at the basal membrane of the cells. The laser spot was focused on the bottom membrane by maximising the fluorescence intensity. To avoid the crosstalk from internalized fluorophores, measurements were done at the bottom of the nucleus (Figure S1). For zebrafish embryos, similarly the laser focus was placed on the membrane by maximizing the fluorescence signal. 3-5 curves were obtained for each spot (five seconds each).

The obtained curves were fit using the freely available FoCuS-point software^66^ using 2D and triplet model.

All scanning FCS experiments were performed on the Zeiss LSM 780 using the 40x 1.2 NA FCS water objective as described previously^43^. TF fluorescence was excited using the 488 Argon laser passing through a 488/594/633 MBS and fluorescence detected using the hybrid GaAsP detector (photon counting mode) in a range from 500–600 nm. All sFCS measurements were obtained as line scans with a length of 5.2 µm (52 pixels with a pixel size of 100 nm) at the bottom membrane of live CHO cells right below the nucleus. Maximum scanning frequency (2081 Hz) was used and data acquired for 50 seconds. The scans were correlated with the FoCuS-scan software^67, 68^. For all data sets the first 10 seconds were cropped off to remove initial bleaching and an 18 second photobleaching correction by local averaging applied. The correlation curves were fitted including 10 times bootstrapping with a 2D one component model (no anomalous subdiffusion, no triplet etc.) and the resulting fitting parameters (transit time, counts per molecule) exported for further analysis.

### Scanning FCS statistical analysis

Statistical analysis of sFCS data was performed as described previously^43^. Briefly, the exported transit times from the FCS fits were histogrammed using Matlab. The resulting transit time histograms were fitted with a lognormal function (for single component/free) or with a double lognormal function (for hindered diffusion). Model selection was performed using maximum likelihood estimation and employing the Bayesian Information criterion (given the data which model works best). To obtain increased fitting accuracy the data were cumulatively, linearly and logarithmically histogrammed and fitted with the respective lognormal function. Thus a large data set of >10 cells with multiple measurements each resulting in > 1000 curves for one repetition can be summarized in the statistical analysis fitting parameters.

### % Lo calculation

ImageJ-Line profile was used to calculate the Lo % as described in refs^14^. A line was selected which crosses the opposite sides of the equatorial plane of the GPMVs having different phases on opposite sides. Opposite sides are chosen to eliminate the laser excitation polarization artefacts. Then, %Lo was calculated as;

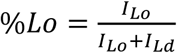

where I is the fluorescence emission intensity. If *%Lo* > 50 %, a lipid analog prefers the liquid ordered phase.

### Molecular Simulations of DPPC:DOPC:Chol membranes

240 DOPC, 116 DPPC, and 44 CHOL per leaflet were assembled into a bilayer and solvated with 40 TIP3P waters using the CHARMM-GUI^69^ membrane builder. The system was equilibrated at a constant temperature of 298 K and constant pressure of 1 bar using semi-isotropic pressure coupling with NAMD v 2.7 on local resources for 50 nsec. The system was then transferred to the Anton special purpose supercomputer^70^ for production simulation. The equations of motion were integrated with the Verlet algorithm with a time step of 2.0 fs. A constant temperature and a pressure of 1 atm were maintained by the Martyna− Tobias− Klein^71^ method, with the pressure coupling effected every 240 fs and the temperature coupling every 24 fs. Lennard-Jones interactions were truncated at 10.14 Å by a hard cutoff with no shift. Long range electrostatics were computed by the k –space Gaussian split Ewald method^72^ on a 64 Å∼ 64 Å∼ 64 point grid, with the parameters of the Gaussian chosen to yield a root mean squared error in the electrostatic force calculation of 0.18%. The total duration of the production simulation was 1.2 μsec. Full simulation details can be found in ref^73^.

### Monte Carlo Simulations of Diffusion Modes

For the simulation of free and hindered diffusion we used and slightly modified the nanosimpy python repository as described previously. 350 particles were randomly initiated and moved at every time step (1 µs) according to having a diffusion coefficient of 1 µm^2^/s. Diffusion was simulated for 20 seconds on a circle with a diameter of 3 µm (wrapped around on the edges). A FCS observation spot (approximated by a Gaussian) was placed in the centre of the simulation area and the apparent intensity sampled at every time step. The observation size was tuned from 250 to 50 nm (STED-FCS diffusion law measurement) to make sure the respective diffusion modes were obtained. For trapped diffusion we employed a statistical model for molecular complex formation rending a random particle immobile for a short time (p_trap_ = p_untrap_ = 0.00005). For hop diffusion we generated a Voronoi mesh with a characteristic mesh size (given as 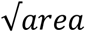) of 110 nm. When a particle would hit a boundary it only had a low chance (phop = 0.05) to pass on to the next compartment and otherwise bounce off. FCS curves were fitted with the FoCuS_point software using a simple 2D diffusion model^66^. Note that for the case of hindered diffusion modes anomalous sub-diffusion can occur (meaning the α-value can drop below 1, ranging from 0.65 −1).

## Acknowledgement

We thank the Wolfson Imaging Centre Oxford and the Micron Advanced Bioimaging Unit (Wellcome Trust Strategic Award 091911) for providing microscope facility and financial support. We acknowledge funding by the Wolfson Foundation, the Medical Research Council (MRC, grant number MC_UU_12010/unit programmes G0902418 and MC_UU_12025), MRC/BBSRC/EPSRC (grant number MR/K01577X/1), the Wellcome Trust (grant ref 104924/14/Z/14), the Deutsche Forschungsgemeinschaft (Research unit 1905 “Structure and function of the peroxisomal translocon”), Oxford-internal funds (John Fell Fund and EPA Cephalosporin Fund) and Wellcome Institutional Strategic Support Fund (ISSF). ES is funded by the Newton-Katip Celebi Institutional Links grant (352333122).

