## Supplemental Figures for "Nanoscale dynamics of cholesterol in the cell membrane"

### Supplementary Information

#### Supplementary Figures

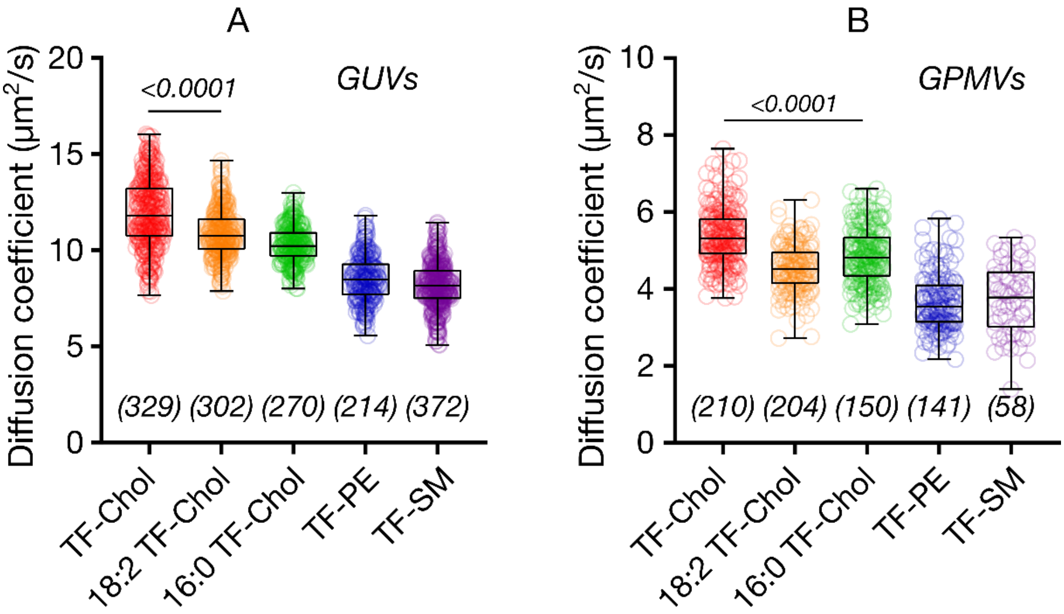


**Figure S1.** Diffusion of cholesterol analogue TF-Chol compared to phospholipid and sphingolipid analogues in model membranes in A) GUVs and B) GPMVs. Data is pooled from at least three independent measurements. Data are shown as box-and-whisker plot showing median, first and third quartiles, and all the data points. Number of data points are indicated on the graphs.

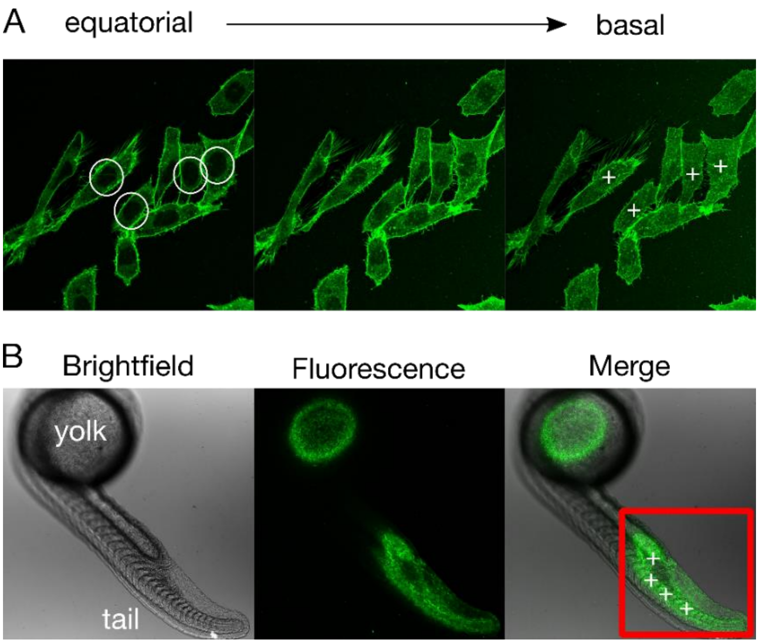


**Figure S2.** FCS measurements in A) live cells and B) embryos labelled with TF-PE. A) To avoid background, we performed the FCS measurements on the bottom membrane underneath the nucleus where there is no internal background. B) For zebrafish embryos, we measured FCS close to the tail region away from the yolk which accumulated lipid analogues.

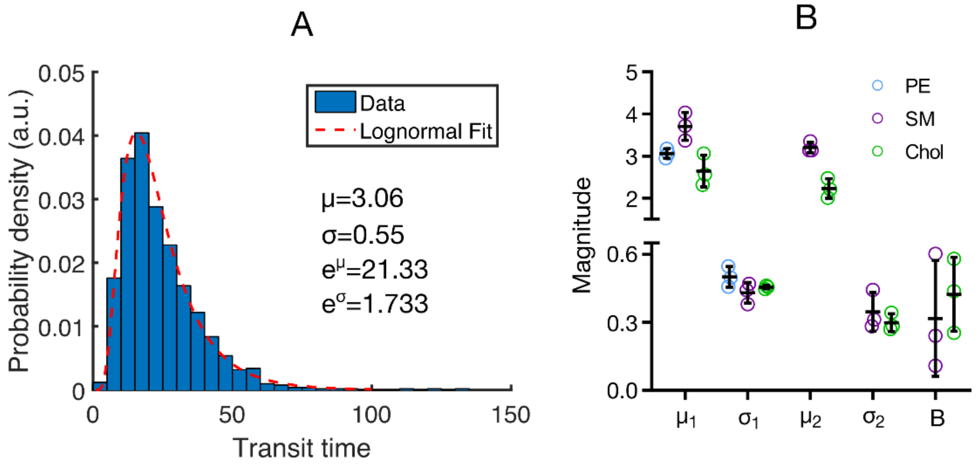


**Figure S3.** A) Simulated lognormal distribution of sFCS transit time data and the respective fitting parameters for free diffusion with an apparent transit time of 21.33 ms. B) sFCS fitting parameters obtained from the lognormal fit for the three fluorescent lipid analogues. TF-PE histograms can be fitted with a single lognormal function whereas a double lognormal fit needs to be employed for TF-SM and TF-Chol.
